# A genome-wide analysis of Cas9 binding specificity using ChIP-seq and targeted sequence capture

**DOI:** 10.1101/005413

**Authors:** Henriette O’Geen, Isabelle M. Henry, Mital S. Bhakta, Joshua F. Meckler, David J. Segal

## Abstract

Clustered regularly interspaced short palindromic repeat (CRISPR) RNA-guided nucleases have gathered considerable excitement as a tool for genome engineering. However, questions remain about the specificity of their target site recognition. Most previous studies have examined predicted off-target binding sites that differ from the perfect target site by one to four mismatches, which represent only a subset of genomic regions. Here, we use ChIP-seq to examine genome-wide CRISPR binding specificity at gRNA-specific and gRNA-independent sites. For two guide RNAs targeting the murine *Snurf* gene promoter, we observed very high binding specificity at the intended target site while off-target binding was observed at 2- to 6-fold lower intensities. We also identified significant gRNA-independent off-target binding. Interestingly, we found that these regions are highly enriched in the PAM site, a sequence required for target site recognition by CRISPR. To determine the relationship between Cas9 binding and endonuclease activity, we used targeted sequence capture as a high-throughput approach to survey a large number of the potential off-target sites identified by ChIP-seq or computational prediction. A high frequency of indels was observed at both target sites and one off-target site, while no cleavage activity could be detected at other ChIP-bound regions. Our data is consistent with recent finding that most interactions between the CRISPR nuclease complex and genomic PAM sites are transient and do not lead to DNA cleavage. The interactions are stabilized by gRNAs with good matches to the target sequence adjacent to the PAM site, resulting in target cleavage activity.

## INTRODUCTION

Targeted genome engineering by nucleases has enabled researchers to alter genetic content in a variety of cell types and organisms. In particular, the RNA-guided Cas9 endonuclease, adapted from the CRISPR system of *Streptococcus pyogenes*, has emerged as the universal tool of choice for advancing biological research as well as the potential for therapy of genetic diseases (Mali et al. 2013b; Sander and Joung 2014). Cas9 is guided to genomic loci by the guide-RNA (gRNA) scaffold containing 20 nucleotides complementary to the genomic target site, which is immediately upstream of a protospacer adjacent motif (PAM) site. The PAM site consists of three nucleotides NGG and is a requirement for Cas9 binding to its target region (Jinek et al. 2012; Sternberg et al. 2014).

Cas9 and its derivatives offer a repertoire of functions. Wild-type Cas9 nuclease acts by introducing double strand breaks at the DNA target site that are either repaired by NHEJ (non homologous end joining) or HR (homologous recombination). Unless donor DNA is provided, the default pathway of NHEJ causes insertion or deletion mutations (indels) (Jinek et al. 2012; Mali et al. 2013c). In addition, Cas9 can be converted to a nickase by a single D10A mutation or to a nuclease-inactive DNA binding protein (dCas9) by introducing two amino acid changes (D10A and H840A) in the RuvC1 and HNH-like nuclease domains, respectively (Jinek et al. 2012). The Cas9 double mutant dCas9 can be fused to a heterologous effector domains to regulate transcription (Gilbert et al. 2013; Maeder et al. 2013; Perez-Pinera et al. 2013). In addition, Cas9 can be fused to domains that regulate the epigenetic landscape at endogenous loci. This strategy has shown potential for zinc finger and TALE DNA binding proteins (Stolzenburg et al. 2012; Konermann et al. 2013; Mendenhall et al. 2013; Johnson et al. 2014). The versatility and ease of use make CRISPR/Cas9 a powerful tool for genome editing and gene regulation, but our understanding of binding specificity and target recognition remains limited.

Recent studies have identified indels introduced by Cas9 nucleases at off-target sites that share sequence similarity to the target site (Cradick et al. 2013; Fu et al. 2013; Hsu et al. 2013; Pattanayak et al. 2013; Cho et al. 2014). All studies were in concurrence about the importance of the PAM site and concluded that mismatches in the 5’ region of the target site were much better tolerated than in the PAM-adjacent sequence, also referred to as the seed region. The seed region has been defined as the sequence of 6 to 12-bp immediately upstream of to the PAM site. However, the search for off-target events was limited to predicted off-target sites, and thus subject to potential biases based on the quality of the predictions. In some studies, Cas9 specificity was assayed by allowing mismatches in the gRNA complementary to the target site. While many of these sites showed off-target activity *in vitro*, specificity varied greatly at genomic sites. For example, Pattayanak *et al* identified 32 potential off-target sites *in vitro*, but only three displayed above-background activity in their genomic context (Pattanayak et al. 2013). Other studies determined off-target effects at endogenous loci that differed by one to six positions from the on-target sequence (Fu et al. 2013; Hsu et al. 2013; Cho et al. 2014). However, it has become clear that off-target effects vary greatly by site, and it has proven difficult to identify a pattern to predict binding specificity. A less biased method, whole-exome sequencing of individual clones isolated after CRISPR/Cas9 nuclease treatment, did not identify any off-target events (Cho et al. 2014). However, this analysis examined 2% of the genome representing only coding regions. Whole genome sequencing of treated pools of cells would provide a better assay but sequencing depth requirements make it unfeasible for efficient detection of off-target effects.

Target site recognition and DNA binding are required prior to CRISPR/Cas9 activity. ChIP-seq (Chromatin Immunoprecipitation followed by high throughput sequencing) is a powerful tool to identify genome-wide binding sites in an unbiased manner. Recently, ChIP-seq has been used to determine binding of CRISPR/Cas9 using four different guide RNAs in mouse stem cells (Wu et al. 2014). The number of off-target sites varied with the amount of expressed Cas9 protein and was also dependent on gRNA target sequence. Different gRNA sequences resulted in highly variable numbers of identified off-target binding sites, ranging from 26 to 5,957. Among the 9,594 gRNA-specific peaks, a subset of 295 (3%) Cas9-bound regions was chosen for validation of off-target cleavage activity using targeted PCR and sequencing. Cas9 cleavage activity was only detected at one of the chosen ChIP-bound sites.

Off-target site predictions based on sequence similarity or genome-wide binding analysis by ChIP-seq have not been reliable predictors of off-target cleavage by CRISPR/Cas9 (Cradick et al. 2013; Fu et al. 2013; Hsu 2013; Mali et al. 2013a; Pattanayak et al. 2013; Cho et al. 2014; Sander and Joung 2014; Wu et al. 2014). Validation of off-target cleavage activity has largely been based on a selection of genomic off-targets that were PCR amplified and then screened for indels using either high throughput sequencing or Sanger sequencing. Clearly, there is still a need for a high-throughput method that allows simultaneous validation of a large number of potential off-target sites.

Here, we employed ChIP-seq analysis followed by a targeted sequence capture approach to determine RNA-guided dCas9 specificity and off-target endonuclease activity on a genome-wide scale. We used two replicates for two different gRNA targets, which allowed us to identify gRNA-specific binding as well as binding events that occur independently of the gRNA-specified target sequence. Since neither ChIP-seq binding, nor off-target site prediction based on sequence similarity alone are good predictors for off-target cleavage activity, there is a need for a comprehensive approach to validate a large number of potential off-target sites. We have demonstrated that sequence capture can be used as an efficient approach to interrogate Cas9 nuclease off-target activity at a large number of genomic regions simultaneously. This approach allowed us to determine and compare cleavage activity at binding sites identified by both ChIP-seq and computationally predicted off-target sites.

## RESULTS

### Genome-wide analysis reveals Cas9 on- and off-target binding with strongest affinity for the perfect target site

ChIP assays are most commonly used to detect protein-DNA interactions in cells. In the case of the Cas9:gRNA:DNA complex, we are interested in protein-DNA interactions that are facilitated by the guide RNA. Active Cas9 nuclease was not used in ChIP assays since it induces indel mutations at cleavage sites which would interfere with Cas9 binding (Jinek et al. 2012). Therefore, we used a D10A and H840A catalytically inactive Cas9. To create a Flag-tagged dCas9 DNA-binding protein (Fig. 1A, top), we fused a short linker (GGGGS) and a 3X Flag tag to the C-terminus of the nuclease-inactive, human codon-optimized Cas9 protein (Mali et al. 2013b).

**Figure 1.**
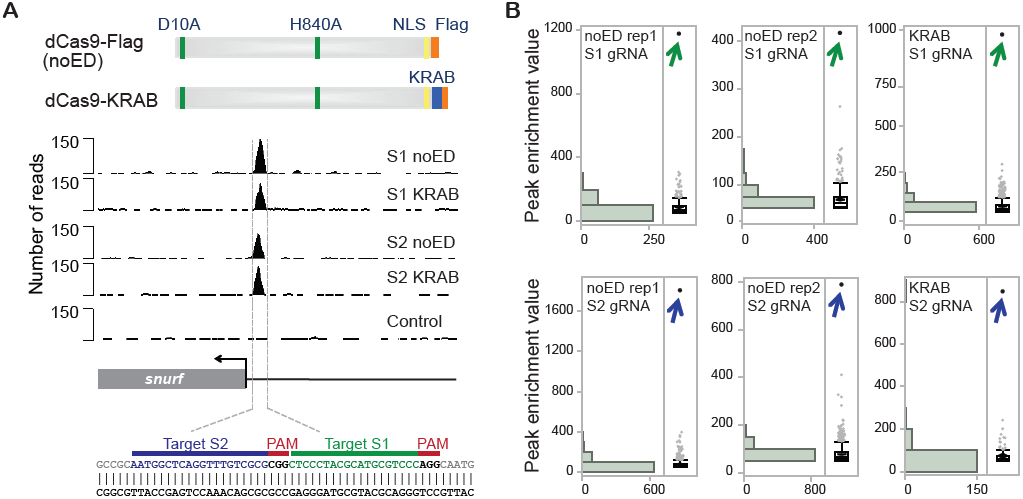
Genome-wide analysis of RNA-guided dCas9 binding in Neuro-2a cells. **(A)** Diagram of the Flag-tagged Cas9 double mutant (dCas9) used in ChIP-seq assay. dCas9 without an effector domain (dCas9-Flag) or fused to a KRAB repressor domain (dCas9-KRAB) are shown at top. Location of nuclear localization domain (NLS) is indicated. ChIP-seq profile of dCas9 binding at the on-target site in the mouse *Snurf* locus is shown in the middle. Binding is shown for dCas9-Flag without an effector domain (noED) or with the KRAB repressor domain (KRAB). RNA-guided binding via S1 and S2 gRNAs is depicted. A U6 promoter plasmid that did not express functional gRNA was used as a control. Nucleotide sequence targeted by S1 and S2 gRNA is shown at bottom. **(B)** Distribution of ChIP-seq peak enrichment is shown for two experiments (rep1 and rep2) for dCas9-Flag without an effector domain and for dCas9 fused to the KRAB repressor domain. ChIP-seq enrichment values were obtained using the MACS1.4 peak caller using genomic input DNA as a control. Results are shown for S1 and S2 gRNAs. On-target binding to the *Snurf* promoter is indicated with an arrow.

Two gRNAs were designed to target two adjacent loci (S1 and S2) within the *Snurf* gene promoter (Fig. 1A, bottom). S1 and S2 gRNAs were carefully selected for their uniqueness in the genome. The closest match to the S1 and S2 target sites in the mouse genome contained 3 mismatches. The mouse neuroblastoma cell line Neuro-2a was cotransfected with a plasmid expressing the dCas9-Flag protein and another expressing one of the two gRNAs. ChIP-seq experiments were carried out twice for each gRNA. Experiment 1 was transfected with 1 μg of dCas9-Flag plasmid and harvested after 24 hours, while experiment 2 was transfected with 7.5 μg of dCas9-Flag plasmid and harvested after 48 hours. The amount of gRNA-expressing plasmid was kept constant at 7.5 μg.

Genome-wide binding sites were identified using the MACS1.4 peak caller using non-enriched chromatin DNA (also referred to as input) as background (Zhang et al. 2008). To reduce the occurrence of false positive peaks, repeat-masked regions were excluded from ChIP-seq analysis (Pickrell et al. 2011). To identify unspecific binding regions, ChIP-seq analysis was performed in cells treated with dCas9-Flag and a control U6 promoter plasmid that did not express a functional gRNA. One unspecific peak was identified in this control data set, which was removed from all other data sets. ChIP-seq analysis of dCas9-Flag binding guided by S1 gRNA resulted in 338 and 517 peaks, while S2 gRNA guided binding resulted in 737 and 1009 peaks (experiments 1 and 2, respectively). The presence of more dCas9-Flag protein in experiment 2 correlates with an increase in ChIP-seq binding sites.

For both gRNAs, the top-ranked binding site in each data set was an identical match to the target locus, demonstrating high affinity for the on-target site (Fig. 1A, middle and Fig. 1B). In experiment 1, for which lower amounts of dCas9-Flag had been used, the target site was enriched 4 and 6 fold higher than the highest off-target binding site for S1 and S2, respectively. The target site enrichment was only two fold higher than the highest off-target site in experiment 2, consistent with the idea that reducing protein and/or gRNA amounts and exposure time can limit off-target binding (Pattanayak et al. 2013; Sternberg et al. 2014; Wu et al. 2014).

### ChIP-seq identifies gRNA-specific and gRNA-independent binding sites

We also wanted to explore the possibility that some of the off-target binding sites represented dCas9 binding events that were not specific to a given gRNA or target sequence. To identify possible sites of gRNA-independent recognition, we searched for peaks that occurred at least once in the S1 gRNA and once in the S2 gRNA data sets and identified 150 binding sites that were not specific to a single gRNA. These gRNA-independent peaks were subtracted from the S1 and S2 data sets and were analyzed separately. Consequently, 274 and 404 peaks specific to S1 gRNA, and 665 and 883 peaks specific to S2 gRNA remained (experiment 1 and experiment 2, respectively). We next overlapped gRNA-specific peaks from the two experiments to obtain 69 S1-specific and 254 S2-specific peaks *bona fide* binding sites (Fig. 2A).

**Figure 2.**
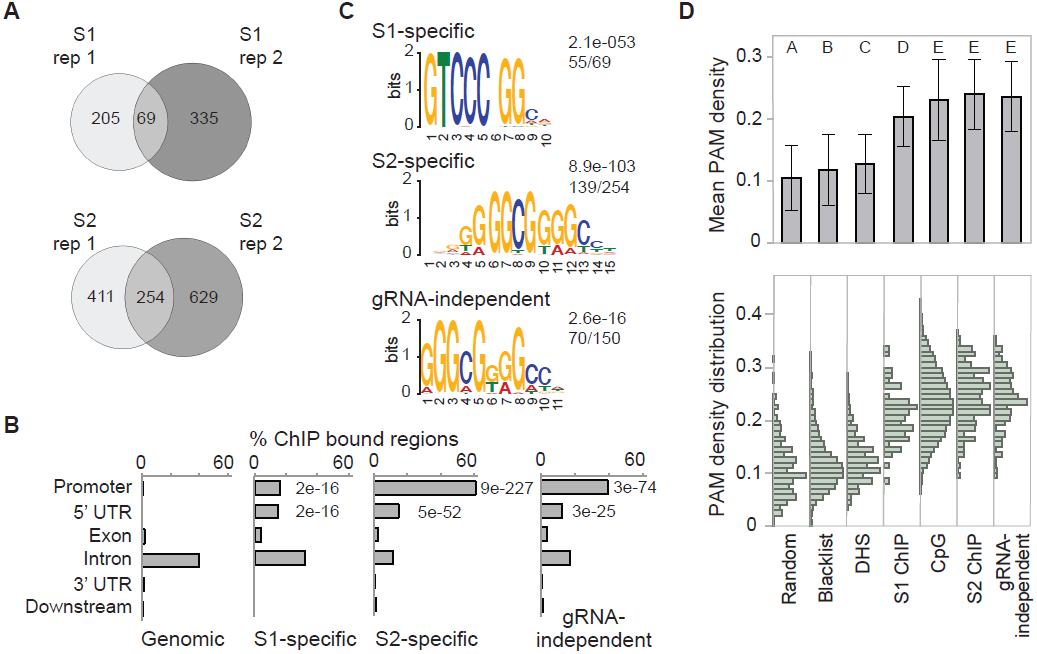
Identification of G- and C-rich motifs at off-target binding sites correlates with increased PAM motif density. **(A)** Venn diagrams show the overlap between ChIP-seq peaks from two replicates. gRNA-independent peaks that are present in S1 and S2 gRNA data sets are referred to as gRNA-independent peaks and were subtracted from S1 and S2 gRNA-specific binding peaks prior to overlap analysis. **(B)** Location analysis of ChIP-seq binding sites. ChIP peaks at gene-proximal regions and within gene bodies were analyzed using Cis-regulatory Element Annotation System (CEAS). Gene-proximal regions include promoters up to 1 kb upstream of the transcription start site (TSS) and downstream regions up to 1 kb downstream of the transcription termination site. Binding within gene bodies is divided into four functional categories: 5’UTRs, exons, introns and 3’UTRs. Localization of ChIP regions within these categories is compared with the genome background percentages for the same categories. P-values are indicated for categories with significant enrichment in ChIP samples as compared to genomic background. **(C)** Identification of *de novo* motifs for overlapping dCas9 binding sites in Neuro-2a cells. MEME-ChIP motif analysis using the central 100 bp of gRNA-specific and gRNA-independent ChIP-seq binding regions reveals the indicated motif. The number of sequences containing the identified motif relative to the total number of binding sites is indicated. **(D)** PAM density plot based on occurrence of NGG in the central 100 bp of gRNA-specific and gRNA-independent ChIP-seq peaks. Both DNA strands were scanned for the three-nucleotide motif. Randomly selected 100-bp regions of the mouse genome and an array of artifact peaks (also known as ENCODE ChIP-seq blacklist) were used as control sets. PAM density was also elevated in CpG islands. Median of PAM density and standard deviation is shown in the top half, while PAM density distribution is plotted in the lower half. Statistical groupings A-E as determined by student T test are indicated above each column.

To determine functional genomic regions bound by Cas9, we used the Cis-regulatory Element Annotation System CEAS (Ji et al. 2006; Shin et al. 2009). Gene-proximal regions include gene promoters up to 1 kb upstream of the transcription start site (TSS), gene bodies and downstream regions up to 1 kb downstream of the transcription termination site (Fig. 2B, Table S1). The majority of gRNA-specific as well as gRNA-independent ChIP peaks localize to gene-proximal regions with 70%, 92% and 74% of S1-specific, S2-specific and gRNA-independent binding sites, respectively. Functional regions of gene bodies include 5’ UTRs, coding exons, introns and 3’UTRs. Localization of ChIP regions within these categories is compared with the genome background percentages for the same categories and *p*-values are calculated. ChIP peaks show significant enrichment at proximal promoters and in 5’UTRs, while less than 5% of exons and less than 1% of 3’UTRs were targeted (Fig. 2B). Binding to intronic regions was not significantly enriched when compared to genomic background.

### The KRAB effector domain does not contribute to dCas9:gRNA recruitment to DNA

The nuclease-inactive dCas9 can be used as a transcriptional regulator when fused to an effector domain, which can either up or down regulate transcription (Gilbert et al. 2013; Maeder et al. 2013; Perez-Pinera et al. 2013) or alter the epigenetic landscape surrounding the target site. Our study was carried out with dCas9-Flag that does not contain an effector domain (noED) to identify direct interactions with genomic DNA. But it has been of concern that effector domains themselves could contribute to the recruitment of fusion proteins to DNA resulting in additional set of off-target binding sites specific to the effector domain. To test this hypothesis, we used the dCas9 protein fused to the KRAB repressor domain and 3X Flag tag (Fig. 1A, top). ChIP-seq experiments were conducted using dCas9-KRAB with either S1 or S2 guide RNAs. After removal of previously identified gRNA-independent peaks, 617 and 166 binding sites were identified for S1 and S2 gRNA, respectively. Genome-wide binding profiles using the dCas9-KRAB fusion protein showed the same distribution as the ones that did not contain an effector domain. The on-target site was by far the most highly enriched genomic region and was at least three fold higher than any off-target binding sites (Fig. 1A, middle and Fig. 1B). We then compared ChIP peaks bound by dCas9-KRAB to the high-confidence ChIP peaks bound by dCas9 without the effector domain. When comparing S1 peaks, 83% of 69 S1-specific binding sites were also occupied by dCas9-KRAB. For S2-specific binding, 57 of the 136 regions bound by dCas9-KRAB were previously identified high-confidence S2 peaks. The lack of highly enriched ChIP peaks in addition to the target site indicates that the KRAB effector domain was not responsible for detectable recruitment to genomic loci. Future studies will elucidate if this holds true for other effector domains such as activators and epigenetic modulators.

### CRISPR/Cas9 complex binds to a 5-bp core seed region and GC-rich genomic regions

The binding profiles for both gRNAs revealed that enrichment values for the target site were by far the highest signal genome-wide (Fig. 1B). The lower enriched off-target binding sites may reflect transient interactions between dCas9 and the genome or may be stable interactions only occurring in a few cells at a time. To identify common features of ChIP binding sites, we performed *de novo* motif analysis on the ChIP peaks identified. We retrieved 100-bp sequences centered around the peak middle of Cas9-bound regions and identified the most significant motif using MEME-ChIP (Machanick and Bailey 2011) (Fig. 2C). 55 out of 69 S1 binding sites contained a motif of <u>GTCCC</u>NGG (E-value 2.1e-53). Interestingly, this motif is identical to the 5-bp sequence immediately upstream of the PAM site (NGG) present in the target site sequence CTCCCTACGCATGC<u>GTCCC</u>(AGG) (Fig. 1A, bottom). Most studies identified a longer 12-bp seed region (Fu et al. 2013; Hsu et al. 2013; Sternberg et al. 2014). We therefore defined the 5-bp directly adjacent to the PAM as the core region. Motif analysis of the S2-bound regions identified a gGGCGgggc motif in 139 of 254 binding sites (E-value 8.9e-103). There is no obvious resemblance between the identified motif and the 5-bp core region of the S2 target site sequence AATGGCTCAGGTTTG<u>TCGCG</u>(CGG). Moreover, it is almost identical to the gGGcGggGc also found in 70 of 150 gRNA-independent binding sites (E-value 2.6e-16). The high percentage of G and C nucleotides is striking in all identified motifs. No motif was identified when a set of randomly chosen genomic regions was analyzed.

We next wanted to interrogate Cas9-bound regions for the presence of partial matches to the target site. It has been shown that the seed region, *i.e.*, the region directly upstream of the PAM site, is most important for target site recognition. We first looked for sequence similarity allowing up to four mismatches to the 12-bp seed region in S1 ChIP-seq peaks, more specifically in 100-bp sequences centered on the middle of ChIP-seq peaks. Among 68 S1 off-target binding sites, 20 sites (29%) contained a 12-bp sequence with up to four mismatches adjacent to a PAM; two sites had two mismatches, four sites had three mismatches and the rest had four mismatches (Supplemental Table S2). When Cas9-bound sites were interrogated for presence of the 5-bp core region adjacent to a PAM, 51 (75%) of the 68 S1 off-target regions contained a perfect match to GTCCC(NGG). A second PAM of NAG has been reported as an alternative to NGG (Hsu et al. 2013). When searching the 68 S1 off-target regions for a perfect match to the sequence GTCCC(NAG), we identified only two sites, suggesting that the PAM motif NAG does not play a major role in target recognition.

We then performed the same analysis for S2 gRNA-specific ChIP bound regions. When allowing for four mismatches, only 22 of the 253 off-target sites contained a sequence similar to the 12-bp seed region adjacent to the PAM motif (NGG). On the other hand, 12 binding sites matched perfectly the 5-bp core region adjacent to the PAM motif, TCGCG(NGG), and another 106 sites exhibited only one mismatch to the core region. The fact that only few binding sites match the 5-bp core region adjacent to a PAM and that the motif for S2 and gRNA-independent binding sites are identical suggests that a subset of S2-specific binding sites might be gRNA-independent binding sites, which we were unable to identify.

### The PAM NGG is overrepresented in Cas9:gRNA bound regions

To investigate the correlation between GC-rich regions bound by Cas9:gRNA and the occurrence of NGG PAM sites, we counted the frequency of PAM sites (NGG) in 100-bp sequences centered on the middle of each ChIP-seq peak. PAM site frequencies were recorded for both DNA strands. The median PAM densities were higher (20% at S1-specific peaks and 24% for S2-specific and gRNA-independent peaks) in off-target binding regions compared to random genomic regions (median of 9%) (Fig. 2D). We also compared our binding sites to three other sets of sequences: i) Blacklist regions of the mouse genome were previously identified as signal artifacts in next generation sequencing experiments, independent of cell line and experiment (https://sites.google.com/site/anshulkundaje/projects/blacklists). There was no overlap between ChIP-seq peaks in our study and blacklist regions identified as part of the ENCODE project (Bernstein et al. 2012) and the density of PAM sites in this list of sites was 11%. ii) Recently, a correlation between regions of DNaseI hypersensitivity and off-target binding was reported (Wu et al. 2014). Because there are no DHS data available for the Neuro-2a cell line used in this study, DNaseI hypersensitive sites from two brain data sets (adult and embryonic) were combined instead. In those, PAM motif density was 12%. iii) CpG islands had a PAM density of 24%. Student’s T tests (p < 0.0001) were conducted to compare PAM densities between different sequence data sets. There was no significant difference in PAM density between CpG islands, S2-specific peaks and gRNA-independent peaks (Fig. 2D). All peaks sets and CpG islands were significantly enriched in PAM sites compared to the other three categories (the ENCODE blacklist, the DNaseI hypersensitive sites and the random set).

### Off-target binding correlates with accessible chromatin and GC-skewed genomic regions

Chromatin accessibility as assayed by DNaseI hypersensitivity had been found to be a strong indicator for *in vivo* Cas9 binding (Wu et al. 2014). We therefore overlapped off-target binding sites identified by ChIP-seq with DNaseI hypersensitive sites from mouse brain available from ENCODE (Stamatoyannopoulos et al. 2012). Indeed 85% of S1, 96% of S2 and 92% of g-RNA independent off-target binding sites localize to accessible chromatin regions, confirming a strong positive correlation between accessible chromatin and off-target binding.

Our study and the work of others suggest that chromatin accessibility plays a critical role in Cas9 binding to off-target sites. While Cas9 cleavage activity was reported at methylated target sites (Hsu 2013), a genome-wide study of Cas9 binding has reported a negative correlation between off-target binding and DNA methylation (Wu et al. 2014). It has been proposed that Cas9 binding to the PAM site triggers local unwinding of DNA, which allows the gRNA molecule to pair with one complementary strand of double stranded DNA, forming an R-loop (Jinek et al. 2012; Sternberg et al. 2014). In mammalian genomes, R-loop formation has been implicated in the protection of CpG island promoters from DNA methylation (Ginno et al. 2012). Regions with GC skew displaying strand symmetry in the distribution of G and C nucleotides are an attribute of human unmethylated CpG island promoters (CGI) and are used as a predictive feature of R-loop formation (Ginno et al. 2012; Ginno et al. 2013).

GC skew is a DNA sequence characteristic measuring the strand bias in the distribution of G and C residues. To investigate if there was a correlation between off-target binding sites identified by ChIP-seq and genomic regions with GC skew, regions of GC skew were identified in the mouse genome using the SkewR algorithm as previously described (Ginno et al. 2012). 43%, 89% and 70% of S1, S2 and gRNA-independent off-target binding sites localized to GC skew regions, respectively. Since GC skew is predictive of co-transcriptional R-loop formation (Ginno et al. 2012; Ginno et al. 2013), these observations suggest that preferential off-target binding may occur at regions already occupied by RNA:DNA hybrids. GC-skewed R-loop-prone regions are highly enriched in CpG islands, 5’-UTRs and DNaseI accessible regions which are features associated with off-target binding sites in this study (Figure 2) and by another group (Wu et al. 2014). Interestingly, the motifs identified as highly enriched in the S2-specific and gRNA-independent data sets (Figure 2C) were highly GC-skewed and each carried two consecutive clusters of at least three guanines, making these motifs ideal candidates for initiating R-loop formation (Roy and Lieber 2009). At the same time, GC-rich, GC-skewed regions such as CpG islands show elevated PAM motif density (Figure 2D), making them ideal off-target decoys from a DNA sequence, structural (R-loop), and chromatin viewpoint.

### Cas9-bound regions and sequence-based predicted off-target sites correlate poorly

If sequence similarity to the target site is the main determining factor for off-target binding, one would expect these sites to be bound by Cas9:gRNA in a ChIP-seq experiment. To identify genomic regions that are similar to the target site sequence, we scanned the mouse genome for genomic loci with up to four mismatches to the target site adjacent to a PAM. Since the target regions were carefully chosen for their uniqueness in the genome, 117 and 28 sequences were identified with 3 or 4 mismatches to the target sites S1 and S2, respectively. There were no sequences identified with less than 3 mismatches to the target sites. For a more comprehensive comparison between predicted off-target sites and ChIP-bound regions, we created a union peak file containing ChIP peaks from both biological replicates for each gRNA. Only one predicted off-target site was present in the S2 ChIP-seq union data set and none in the union of S1 peaks. Interestingly, the one site that was in common between S2-specific ChIP bound regions and computationally predicted sites was the same off-target site that contained the best match to the 12-bp seed region (Supplemental Table S2). We labeled this common off-target site as OT1. OT1 was bound with about 8-fold lower affinity in ChIP experiments as compared to the S2 target site. In fact, based on peak enrichment values OT1 was ranked number 10 in experiment 1 and number 140 in experiment 2 for ChIP enrichment in the S2 gRNA specific data set. The low enrichment values suggest that mismatches destabilize the Cas9:gRNA:DNA interaction.

### Sequence capture as a comprehensive method to determine off-target cleavage

To assess Cas9 activity at potential off-target sites, we used sequence capture as a global and efficient approach to simultaneously screen hundreds of potential off-target regions for indels, the hallmark of Cas9 nuclease activity (Gnirke et al. 2009). While exome capture has been used to screen for Cas9 induced indels in transcribed regions of the genome (Cho et al. 2014), custom capture allows us to interrogate very specific loci. Location analysis of ChIP-seq binding sites showed that only 4.4%, 2.8% and 3.4% of binding sites (S1, S2, and gRNA-independent sites, respectively) are found in coding exons (Fig. 2B). Targeted sequence capture allowed us to screen off-target binding sites identified by ChIP-seq as well as computationally predicted genomic regions. Capture baits were designed to cover 200 bp centered on each potential off-target site. Baits were 100 bp in length overlapping by 50 bp. Three baits were designed to cover a 200-bp genomic region.

Custom capture baits were designed to survey a large number of sites including 473 sites bound by ChIP (69 for S1, 254 for S2, 150 for gRNA-independent peaks), 310 computationally predicted sites (170 for S1, 140 for S2) and 430 random control regions. Illumina genomic sequencing libraries were prepared from four samples including untreated Neuro-2a cells, Neuro-2a cells expressing Cas9 endonuclease together with either empty, S1 or S2 gRNA. All four indexed libraries were pooled before sequence capture. Sequence capture was performed and the output was analyzed by paired-end sequencing (100 × 100 nucleotides). Sequence reads were filtered and trimmed based on read quality and aligned to the mm9 mouse genome (UCSC). Identical reads generated by amplification during library preparation were removed from the analysis. To assess the efficiency of the capture reaction, coverage of targeted regions relative to randomly chosen regions was calculated. The mean number of reads mapping to each 200-bp target region ranged from 82 to 111 reads for treated samples and 159 reads for untreated Neuro-2a, while randomly chosen genomic regions showed a mean read count of 0.1 (Supplemental Table S3), which corresponds to a more than 1,000-fold enrichment of captured regions.

To identify indels, we first selected read pairs for which at least one of the two reads unambiguously mapped the captured regions. Of these sequences, we identified read pairs for which the forward and reverse reads overlapped, in which case they were joined to generate one long sequence. These assembled sequence reads were then compared to the target sequence. Reads that were identical matches to the target sequence were identified as wild type (WT). Sequences containing single nucleotide polymorphisms (SNPs) as compared to the reference were recorded, but were not considered a result of Cas9 cleavage activity. Insertions and deletions were identified and the percentage of indels was calculated relative to all reads matching each target region. A minimal threshold of 25 sequence reads per target region was applied for indel analysis. A certain level of background indels has been reported (Pattanayak et al. 2013; Cho et al. 2014; Wu et al. 2014) and is mostly thought to be the result of sequencing errors. Furthermore, it has also been reported that indel rates can increase to about 2% in homopolymer stretches (Minoche et al. 2011). Since we observed an elevated indel frequency next to homopolymers in all samples including untreated cells, we omitted these targeted sequences from analysis. We then compared indels in gRNA-treated samples with indels in control samples and applied the Fisher’s exact test to identify genomic regions that had statistically significant evidence of indels (n = 1265, p < 0.01). Only control samples that had less than 2% indel frequency were compared to samples treated with S1 or S2 gRNA. No indels with statistical significance were identified when the reverse analysis was carried out (Supplemental Fig. S1).

In Neuro-2a cells treated with S1 or S2 gRNAs, the two on-target sites were identified as statistically significant with an indel frequency of 22% and 14.3%, respectively (Fig. 3A, Fig. 3B). While no significant off-target cleavage was observed for S1 gRNA, one off-target site was identified for S2 gRNA. In this experiment, the indel frequency of 19.7% observed at the off-target site (OT1) was higher than 14.3% at the target site (TS) (Fig. 3A, Fig. 3B). No statistically significant site was identified when comparing indel frequency in the control cells using empty gRNA to untreated Neuro-2a (Fig. 3A). The off-target site that was identified by ChIP-seq and validated by sequence capture shows three mismatches to the target sequence. Two mismatches occurred at the very 5’ end, furthest away from the PAM site, while the third mismatch occurred in the 5-bp core region (Fig. 3B). It has been reported that off-target activity is more likely to occur when mismatches between the guide RNA and the template are not adjacent to the PAM site, but rather in the PAM distal region (Fu et al. 2013; Hsu et al. 2013; Pattanayak et al. 2013). However, OT1 also contains a mismatch just two base pairs from the PAM site. OT1 is specific to S2 gRNA in ChIP-seq as well as Cas9 activity assay. No ChIP binding to OT1 was observed using S1 gRNA or control gRNA (Fig. 3C). Even though we found poor correlation between ChIP-bound regions and predicted off-target sites overall, the one site that was identified by both methods (OT1) was also confirmed by the Cas9 cleavage assay. Future work will explore if this holds true for a larger selection of gRNAs.

**Figure 3.**
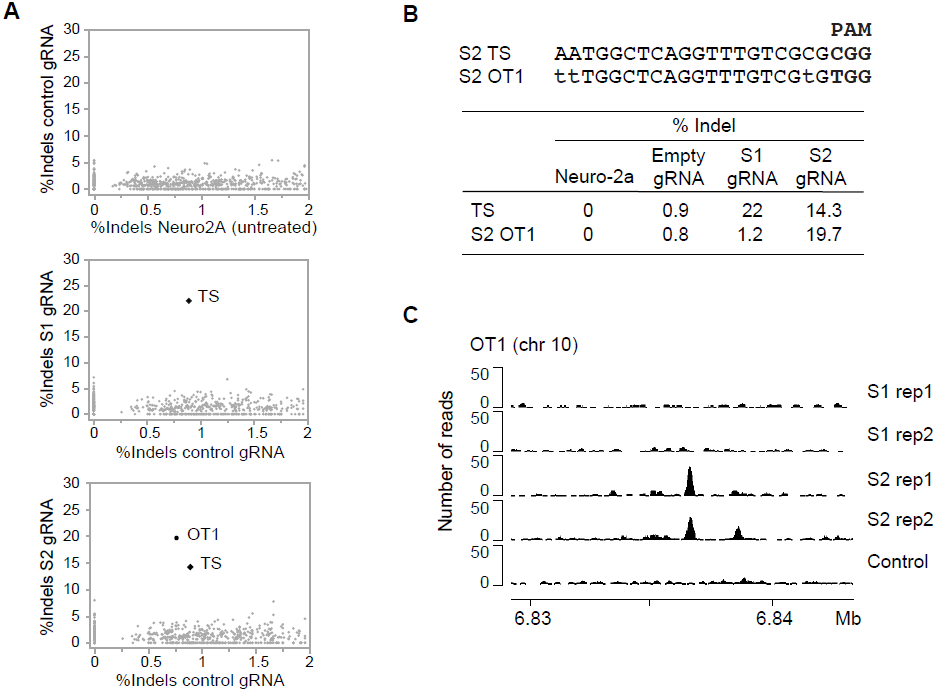
Sequence capture identifies indels at the target sites and one off-target site. **(A)** Bivariate analysis of percent indels relative to the total number of reads per target region. The top panel depicts a comparison of Neuro-2a control cells expressing catalytically active Cas9 and non-functional gRNA to untreated Neuro-2a. The middle panel shows a comparison of cells treated with S1 gRNA against control cells. High occurrence of indels is observed for the target site (TS). The bottom panel shows a comparison of cells treated with S2 gRNA to control cells expressing non-functional gRNA. High indel frequencies are observed for one off-target site (OT1) in addition to the target site (TS). **(B)** A comparison of S2 TS and OT1 sequences, and a table listing the percentage of indels calculated for each of the four data sets at S1 and S2 TS and S2 OT1. **(C)** Browser image depicting the position and height of ChIP-seq peaks to the off-target site OT1 in the mouse genome (mm9 assembly). Binding is specific to cells treated with S2 gRNA and is not present in control cells or cells treated with S1 gRNA.

The majority of indels identified by sequence capture were deletions ranging from 1 to 52 nucleotides (Table 1). For the S1 target site, we identified seven deletion events, one insertion event and two sequences containing both a deletion and an insertion. For the S2 target site, five sequences were identified with deletions and two sequences with an insertion of one nucleotide. The S2 off-target site revealed seven deletion events, four single nucleotide insertion events and one sequence containing a 52-bp deletion in addition to a 2-bp insertion.

**Table 1.**
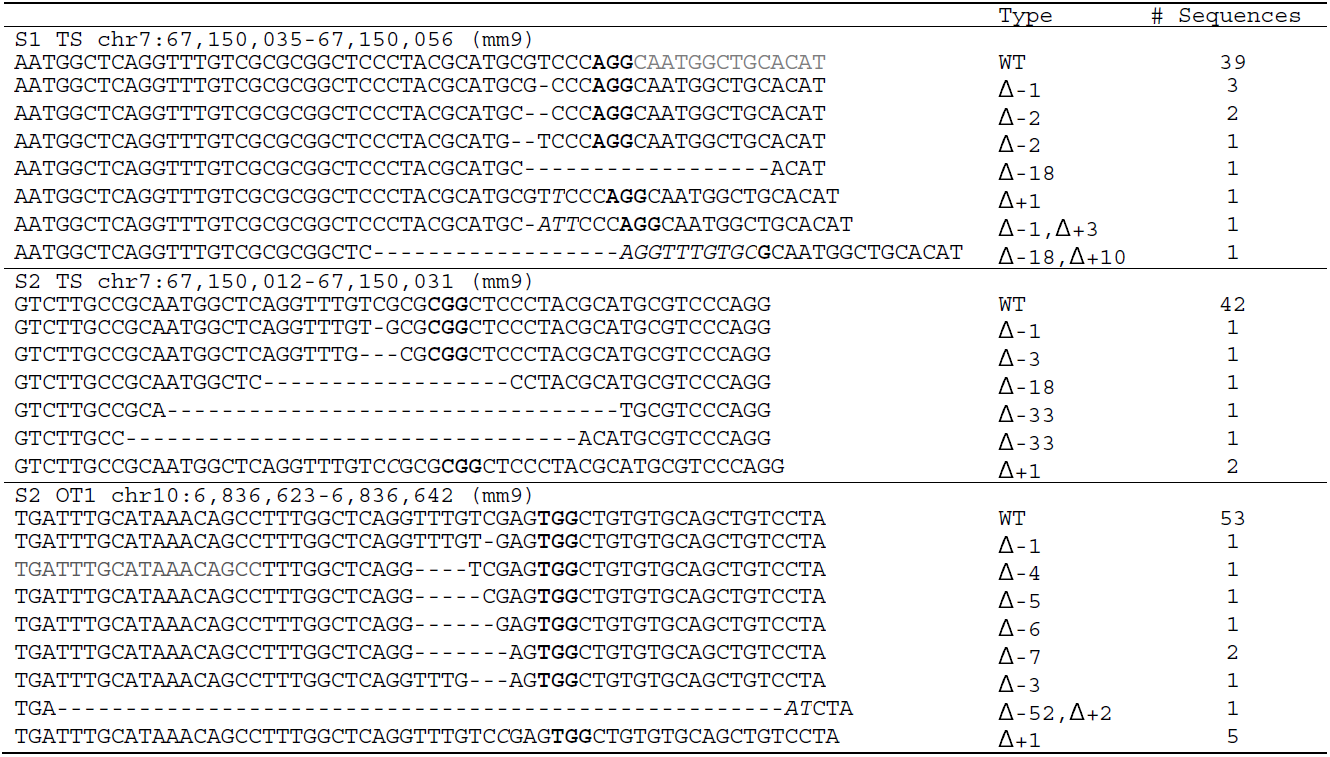
Indels identified by targeted sequence capture at the S1 and S2 target sites (TS) and S2 off-target site (OT1). Wild type sequences (WT) are a perfect match to the target site. Deletions are indicated by dashes and insertions are shown in italic letters. Four sequences had an insertion in addition to the deletion. The three-nucleotide PAM motif is underlined where present. The target sites are shown in black letters while flanking sequences are grey.

## DISCUSSION

We have demonstrated that genome-wide ChIP-seq analysis in combination with targeted sequence capture can be used to assess Cas9 cleavage activity at a large number of potential off-target sites. Custom capture designs are available ranging from 20,000 to 200,000 custom baits, which translates to screening for indels of ∼6,670 to 66,700 genomic loci. This could be very powerful approach to survey off-target activity for a multitude of gRNAs. In our study, we compared indel frequencies at 461 loci identified by ChIP-seq and 310 off-target sites predicted by similarity to target sequence. We confirmed Cas9 cleavage activity at the target sites of both gRNAs and identified a single off-target site (OT1). This off-target site was also the only site that was identified using both the genome-wide ChIP-seq binding analysis and the sequence-based *in silico* prediction. Additionally, we found that binding of Cas9 protein does not necessarily translate into Cas9 nuclease activity. The high enrichment of the target site sets it apart from all off-target sites. This suggests that stable interactions only occur between Cas9:gRNA and the exact target site. Off-target sites that differ from the target site by three or more mismatches do not form as stable interactions, which is reflected in much lower ChIP enrichments and the lack of detectable cleavage activity. OT1 was enriched at much lower levels than the target site in ChIP experiments but cleavage still occurred at high frequency. Our results support previous findings that ChIP enrichment and cleavage activity are not necessarily correlated (Wu et al. 2014). Currently, introduction of indels by Cas9 is used to determine off-target activity. However, the question remains whether dCas9 fusion proteins with e.g. epigenetic modulators display similar binding specificity or perhaps have increased off-target activity.

The uniquely high ChIP enrichment at the endogenous target site observed in our study confirms the high stability and specificity of RNA-guided dCas9 binding in a mammalian genome. Even with increased Cas9 protein amounts, the target site remains by far the most enriched binding site in the genome. Our genome-wide ChIP-seq analysis demonstrates a preference of Cas9:gRNA binding to PAM dense regions in the genome, which suggests that the off-target binding sites represent transient interactions as Cas9 scans along the genome for PAM sites. To our knowledge, this is the first study addressing Cas9:gRNA binding to genomic regions harboring PAM motifs on a genome-wide scale. Our findings are in agreement with target site recognition of CRISPR/Cas9 observed using the 49-kb lambda phage genome (Sternberg et al. 2014). Binding at regions in the lambda DNA genome complementary to gRNA was very stable, while binding to non-target DNA was of transient nature. *In vitro*, these stable on-target binding events could only be displaced with PAM-containing oligonucleotides and were, to some extent, unaffected by the DNA sequence adjacent to the PAM motif. These experiments suggest that Cas9 binding to non-target regions on lambda DNA probably occurred at PAM-rich regions.

In addition to gRNA-specific off-target binding sites, we were able to identify gRNA-independent binding sites that were common to two different gRNAs. gRNA-independent binding sites may represent transient interactions of Cas9:gRNA as it scans along the genome for accessible PAM motifs, while gRNA-specific binding sites use the 5 bp adjacent to the PAM motif as a core seed region. Recently, ChIP-seq analysis identified a 5-bp core region specific to each of the four gRNA sequences used (Wu et al. 2014). It is important to note that our findings using S1 gRNA using dCas9-Flag, which contains a C-terminal fusion of the Flag-tag, concur with the study by Wu et al. (Wu et al. 2014), which used the HA-tag fused to the N-terminus of dCas9. Although, we identified the 5-bp region in the S1 ChIP-seq data set, we were unable to identify a motif with sequence similarity to the target site. The motif identified in the S2 data set was very similar to the one found in gRNA-independent peaks. We therefore speculate that many of these sites are also gRNA-independent that we have not been able to separate yet. Future experiments using additional gRNAs will provide more insight.

A great deal of future research will focus on improving and predicting Cas9 specificity for a given gRNA. It is still unclear why RNA-guided targeting results in high off-target activity for one gRNA, but not for another. It is certainly important to carefully design gRNAs based on the uniqueness of its target sequence in the genome. Careful gRNA design can be combined with other off-target limiting methods. For example, it has been shown that by simply using a shorter complementary gRNA of 18 nucleotides instead of 20 nucleotides, off-target effects became undetectable while on-target activity remained the same (Fu et al. 2014). Another study also reported that shorter gRNA sequences reduced off-target effects, but found that on-target activity was reduced as well (Pattanayak et al. 2013). On the other hand, addition of two G nucleotides at the 5’ end of the gRNA sequence was found to increase target site specificity (Cho et al. 2014). As an alternative to Cas9 nucleases that introduce double strand breaks, the use of paired Cas9 nickases significantly reduced off-target activity (Ran et al. 2013; Cho et al. 2014). However, our results corroborate those of the ChIP-seq study of Wu et al. in suggesting that even a simple configuration of a Cas9:gRNA nuclease can support very specific DNA cleavage activity.

## METHODS

### Cell culture and transfection

The mouse neuroblastoma cell line Neuro-2a (ATCC #CCL-131) was grown in Dulbecco’s modified Eagle’s medium (DMEM) supplemented with 10% bovine calf serum. Neuro-2a cells were grown to 70% confluency and transfected using Lipofectamine 2000 (Life Technologies). For ChIP-seq, cells were transfected with 7.5 μg gRNA-expressing plasmid and 7.5 μg Flag-tagged dCas9 plasmid per 10-cm dish. Cells were cross-linked 24 or 48 hours post transfection by incubation with 1% formaldehyde solution for 10 minutes at room temperature. Cross-linked cell pellets were stored at −80°C.

### Plasmids

Plasmids encoding hCas9-WT and hCas9-D10A were purchased from Addgene (#41815 and #41816, respectively). The double mutant plasmid dCas9 (hCas9-D10A/ H840A) was obtained by site directed mutagenesis. Plasmid dCas9-Flag was generated by adding a 5 amino-acid linker (GGGGS) and 3X Flag tag to the C-terminus of dCas9. Flag-tagged dCas9 plasmid will be made available from Addgene. The KRAB repressor domain was added upstream of the 3X Flag tag by Gibson cloning resulting in plasmid dCas9-KRAB-Flag. The gRNA cloning vector was purchased from Addgene (#41824) and target specific gRNA plasmids were created following recommended guidelines (http://www.addgene.org/static/cms/files/hCRISPR_gRNA_Synthesis.pdf).

### ChIP-seq assay and data analysis

ChIP assays were performed as previously described with minor modifications (O’Geen et al. 2010). 50 μg of sonicated chromatin was incubated with 3 μg of monoclonal anti-Flag antibody (SIGMA M2 F1804). After incubation with 3 μg of rabbit anti mouse serum, StaphA cells (Sigma-Aldrich) were used to collect the immunoprecipitates. After washes and reversal of DNA-RNA-protein cross-links, the entire ChIP sample was used to create an Illumina sequencing library using the KAPA library preparation kit (KAPA Biosystems) and NEXTflex DNA barcodes (BIOO Scientific). Quantitative real-time PCR (qPCR) was performed to confirm enrichment of targets in the ChIP libraries as compared to input libraries. Primers to the mouse *Snurf* gene promoter target site were used as positive control primers (Snurf-F 5’-CTCTCCTCTCTGCGCTAGTC-3’ and Snurf-R 5’-AGAGACCCCTGCATTGCG-3’), while a region on mouse chromosome 4 served as a negative control (mmchr4-F 5’-GAGCTATGGCCCATTGATGT-3’ and mmchr4-R 5’-AATAGTGGGATGGTGGGAGA-3’). Libraries were sequenced using the HiSeq 2000 platform (Illumina). Short sequence reads (SR50) were aligned to the mm9 genome assembly using bowtie2 (Langmead and Salzberg 2012). Binding sites were identified using MACS1.4 with a chromatin input library as the control data set (Zhang et al. 2008). Only binding sites that mapped to the non-repeating sequence of the genome were retained for downstream analysis. Overlap analysis was performed using bedtools.

### Data source

The ENCODE ChIP-seq blacklist was obtained at https://sites.google.com/site/anshulkundaje/projects/blacklists (Bernstein et al. 2012). DNaseI hypersensitivity data (narrow peak file format) from whole mouse brain were downloaded from the UCSC genome browser hosting the mouse ENCODE project (Stamatoyannopoulos et al. 2012). DNaseI hypersensitive sites (DHS) from an adult (week 8) and embryonic (day 14.5) mouse were merged to create a large pool of brain-specific DHS sites. CpG Island coordinates were obtained from the CpGIslandsExt table available at the UCSC Genome browser. All data sets were from the July 2007 mouse assembly (mm9).

### Targeted sequence capture

Neuro-2a cells were co-transfected with equal amounts of plasmids hCas9, gRNA-expression plasmid and pCMV-eGFP. 72 hours after transfection cells were sorted for GFP expressing cells using the Cytomation MoFlo Cell Sorter at the UC Davis Flow Cytometry Shared Resource Laboratory. Genomic DNA was isolated using the Gentra Puregene kit (QIAGEN). Genomic DNA was fragmented to an average size of 150 bp using the BioRuptor NGS (Diagenode). Illumina libraries were prepared with the KAPA library preparation kit (KAPA Biosystems). After ligation to NEXTflex DNA barcodes (BIOO Scientific), DNA was amplified using six PCR cycles following KAPA library preparation kit specifications. Libraries were pooled in equimolar ratios and hybridized to custom designed baits (MYbaits) following the manufacturer’s instructions (MYcroarray). Each targeted sequence was 200 bp long and targeted by three 100-bp MYbait oligonucleotides covering the entire length of the targeted region with a 50 bp walking step. Captured libraries were amplified using 18 PCR cycles. Enrichment of targeted regions was confirmed using qPCR with the same primers that were used for ChIP-seq confirmations. Custom capture libraries were sequenced as paired-ends (PE100) using the Illumina 2500 platform.

### Sequence capture data processing and indel analysis

Sequencing reads were split into individual genomic libraries according to index read sequences using publicly available custom python scripts as previously described ((Henry et al. 2014) and http://comailab.genomecenter.ucdavis.edu/index.php/Barcoded_data_preparation_tools). Briefly, reads were trimmed for quality, the presence of adaptor sequences or N bases and reads that were shorter than 35 bp post-trimming were discarded.

The resulting reads were aligned to the mm9 mouse reference genome using bowtie2 with the *--local* parameter allowing soft clipping of sequence reads. The resulting SAM file, containing information about mapping positions for each read, was screened for the presence of PCR clonal reads as follows: if several read pairs mapped to the same starting positions, and in the same direction, only one of those read pair was retained for downstream analysis. This was performed using a custom python script also available online (overamp.py at http://comailab.genomecenter.ucdavis.edu/index.php/Bwa-doall). The resulting files (non-clonal SAM files) were used for downstream analyses.

Analysis of indel frequencies was performed as follows. First, read pairs for which at least one read mapped at least partially and unambiguously (only one best match found) to the targeted space were selected. Read sequences and associated target peak names were output to a new file. Next, these read-pair sequences were aligned to each other to search for overlap between the two reads. If overlap was found (at least 5-bp overlap), then the two sequences were combined to create a single sequence. If no overlap could be found, the two read pairs were output separately in the same file. Last, the remaining sequences were compared to the reference sequence for each peak to identify polymorphisms and indels. First, the longest region of overlap between the peak and the reference sequence was identified. If the region was smaller than 20 bp, the read was not retained for further analysis. Additionally, the region had to at least span 100 bp. For the adequate regions, region sequences were compared between the read and the reference sequences and reads were divided into four categories: reads that matched the reference sequence exactly were labeled as WT, reads that had the same length but differed from the reference sequence by at least one nucleotide were labeled as “SNP” and reads that were shorter or longer than the reference sequence were labeled as carrying a deletion or an insertion, respectively. For each peak, the percentage of indels was calculated by dividing the number of reads labeled as insertion or deletion by the total number of reads containing adequate regions of overlap with the targeted sequences. Targeted sequences containing homopolymers of more than five nucleotides were removed from analysis.

## DATA ACCESS

All ChIP-seq data have been submitted to the Gene Expression Omnibus and are available under accession number (TBA). Sequence capture data were submitted to the NCBI Short Read Archive (SRA) database (TBA). A website implementing the gRNA selection strategy and Bsite software used to scan the mouse mm9 genome for sequences similar to a given target sequence is available at http://www.genomecenter.ucdavis.edu/segallab/segallabsoftware.

## ACKNOWLEDGEMENTS

We thank members of the DNA Technologies and Expression Analysis Core Facilities of the UC Davis Genome Center for assistance with sequencing and Bridget McLaughlin at the UC Davis Flow-Assisted Cell Sorting (FACS) Cytometry Shared Resource Laboratories for assistance with FACS sorting. We thank Meric Lieberman for advice on data analysis, and Luca Comai and Peggy J. Farnham for helpful discussions. We also want to thank Fred Chedin and Stella Hartono for helpful discussion on R-loop formation and access to mouse GC skew data. This work was funded by grants from the NIH GM097073 (DJS) and HG006761 (PJF), the W.M. Keck Foundation (DJS), and the DOE Office of Science, Biological and Environmental Research (LC and IMH).

## AUTHOR CONTRIBUTIONS

HO and DJS conceived of the study; HO and DJS designed research; HO, MSB, and JFM, contributed materials; HO performed experiments; HO and IMH performed bioinformatics analyses; HO drafted the manuscript; HO, IMH, MSB, JFM, and DJS edited the manuscript. All authors read and approved the final manuscript.

## COMPETING FINANCIAL INTERESTS

The authors declare no competing financial interests.

## Supplementary Data

Supplemental Figure S1. Sequence capture does not identify significant indels in control samples.

**Supplemental Figure S1. Sequence capture does not identify significant indels in control samples. (A)** Bivariate analysis of percent indels relative to the total number of reads per target region. Neuro-2a cells expressing catalytically active Cas9 and one of three gRNA variations are compared. The top panel depicts a comparison of cells expressing non-functional gRNA to cells treated with S1 gRNA, while the bottom compares non-functional gRNA expressing cells against cells treated with S2 gRNA. No significant indels were identified by Fisher’s exact test (n = 1265, p < 0.01). S1 or S2 gRNA-treated samples that had less than 2% indel frequency were compared to non-functional gRNA control sample.

Supplemental Table S1. Location analysis of ChIP-seq peaks at S1, S2 or of gRNA-independent peaks in Neuro-2a cells.

**Supplemental Table S1. Location analysis of ChIP-seq peaks at S1, S2 or of gRNA-independent peaks in Neuro-2a cells.** Chromosomal coordinates of Cas9 bound regions are listed as well as the bp distance to transcription start site (TSS) of the closest RefSeq gene. ChIP-seq peak location within a gene, intronic or exonic localization within that gene is specified. RefSeq gene symbols and chromosomal coordinates of genes are reported.

Supplemental Table S2. Identification of genomic sites with up to four mismatches complimentary to the 12-bp seed region of either S1 or S2 gRNA.

**Supplemental Table S2. Identification of genomic sites with up to four mismatches complimentary to the 12-bp seed region of either S1 or S2 gRNA**. Location analysis was performed to determine bp distance to the closest transcription start site (TSS) of a RefSeq gene.

Supplemental Table S3. Mean coverage over captured regions.

**Supplemental Table S3. Mean coverage over captured regions** The mean number of sequence read mapping to each 200 base pair target region (n = 1,265) is shown for untreated Neuro-2a cells, cells treated with catalytically active Cas9 and non-functional gRNA, S1 gRNA or S2 gRNA. As a comparison coverage was also calculated for a random set of non-targeted genomic regions of the same size (n=1,796). Fold enrichment of capture regions as compared to random regions is shown.

